# Development and validation of 58K SNP-array and high-density linkage map in Nile tilapia (*O. niloticus*)

**DOI:** 10.1101/322826

**Authors:** Rajesh Joshi, Mariann Árnyasi, Sigbjørn Lien, Hans Magnus Gjøen, Alejandro Tola Alvarez, Matthew Kent

**Affiliations:** Department of Animal and Aquacultural Sciences, Faculty of Biosciences, Norwegian University 3 of Life Sciences, N-1432, Ås, Norway; Genomar Genetics AS, Norway

**Keywords:** Tilapia, linkage map, SNP array, genomics, sex determination, amh, anti-Müllerian hormone

## Abstract

Despite being the second most important aquaculture species in the world accounting for 7.4% of global production in 2015, tilapia aquaculture has lacked genomic tools like SNP-arrays and high-density linkage maps to improve selection accuracy and accelerate genetic progress. In this paper we describe the development of a genotyping array containing more than 58,000 SNPs for Nile tilapia (Oreochromis niloticus). SNPs were identified from whole genome resequencing of 32 individuals from the commercial population of the Genomar strain, and selected for the SNP-array based on polymorphic information content and physical distribution across the genome using the Orenil1.1 genome assembly as reference sequence. SNP-performance was evaluated by genotyping 4991 individuals, including 689 offspring belonging to 41 full-sib families, which revealed high-quality genotype data for 43,588 of the SNPs. A preliminary genetic linkage map was constructed using Lepmap2 which in turn was integrated with information from the O_niloticus_UMD1 genome assembly to produce an integrated physical and genetic linkage map comprising 40,186 SNPs distributed across 22 linkage groups. Around one-third of the linkage groups showed a different recombination rate between sexes, with male and female map lengths differing by a factor of 1.2 (1359.6cM and 1632.9cM respectively), with most linkage groups displayed a sigmoid recombination profile. Finally, the sex-determining locus in this population was mapped to position 40.53 cM on linkage group 23, in the vicinity of the anti-Müllerian hormone (amh) gene. These new resources has the potential to greatly influence and improve the genetic gain when applying genomic selection and surpass the difficulties of efficient selection for invasive traits in tilapia.

## 1 Introduction

Nile tilapia (*Oreochromis niloticus*) is an important fresh-water aquaculture species farmed in more than 100 countries including many developing countries where it is an important source of dietary protein (ADB, 2005). Thanks to its fast growth, short generational interval (5 months), relatively small size, adaptability to different environments, and ease to work with, it is also used as a model species for research into fish endocrinology (Seale et al., 2002), physiology (Vilela et al., 2003; Wright and Land, 1998), and evolutionary and developmental biology (Fujimura and Okada, 2007). Nile Tilapia production is supported by more than 20 breeding programs based mainly in South East Asia (Neira, 2010). Most of the commercial and farmed Tilapia strains are derived from the genetically improved farmed tilapia (GIFT) base strain established in the early 1990s (Eknath et al., 1993), among these the Genomar Supreme Tilapia (GST^®^) strain which has undergone more than 25 generations of selection.

So far Tilapia breeding programs have relied on traditional breeding approaches based on easily measurable phenotypes such as weight and length, and have just recently started to implement modern genome-based strategies, such as marker-assisted and genomic selection (Meuwissen et al., 2016). Compared to livestock species, aquaculture has been slower to adopt genome-based selection tools largely due to a lack of genomic resources such as reference genomes, SNP arrays and linkage maps. But in species where genomics has been used to guide selection (e.g. rainbow trout (Gonzalez-Pena et al., 2016; Vallejo et al., 2015), Atlantic salmon (Ødegård et al., 2014; Tsai et al., 2015) and Common carp (Lv et al., 2016)), there are notable success stories related to disease resistance (Correa et al., 2015, 2017; Moen et al., 2015; Vallejo et al., 2017) and carcass quality traits (Gonzalez-Pena et al., 2016).

The first genome assembly for *O. niloticus* (released in 2011; 0renil1.0, and updated to Orenil1.1 at the end of 2012 (NCBI, 2018)) was based on short-read sequencing. A newer assembly (O_niloticus_UMD1) was generated using a combination of novel long-reads (generated using Pacific BioSciences technology) and publicly available Illumina short reads (Conte et al., 2017). Four linkage maps of varying resolution (Guyon et al., 2012; Kocher et al., 1998; Lee et al., 2005; Palaiokostas et al., 2013) have also been published, with the most recent map containing 3,802 markers (Palaiokostas et al., 2013). These maps were constructed using markers found with Restriction-site Associated DNA (RAD) sequencing, microsatellites and/or AFLP markers. The RAD based strategies usually generate a SNP resource of medium density and are highly efficient in species where a reference genome is not available (Robledo et al., 2017). In comparison, a SNP array offers the advantages of increased genotype accuracy of much higher numbers of markers as well as control over the physical distribution of these across the genome (Robledo et al., 2017). In this paper, we report the development of a 58K SNP-array (Onil50) and construction of a high dense linkage map in the commercial strain of Nile Tilapia, Genomar Supreme Tilapia (GST^®^), which is the continuation of the widespread GIFT-strain.

## 2 Materials and Methods

### 2.1 SNP-array (Onil50) development

#### 2.1.1 Whole genome sequencing and SNP-detection

Genomic DNA from 32 individuals (13 males and 19 females) was extracted from fin-clips (preserved in Ethanol) using Qiagen DNeasy columns (Qiagen, Germany). DNA quality was assessed by agarose gel electrophoresis and quantified using a Qubit fluorimeter (ThermoScientific, USA). After normalization, sequencing libraries were prepared and barcoded using TruSeq sample preparation kit and sequenced (2 × 125) across 10 lanes on an Illumina HiSeq 2500 (Illumina, USA) by a commercial provider. At the time this work was carried out, Orenil 1.1 Tilapia represented the highest quality reference genome available (NCBI Assembly *Oreochromis niloticus*: GCF_000188235.2_Orenil1.1_genomic), and reads were aligned to it using BWA 0.7.12 (Li, 2013) with default parameters. Putative SNPs were identified using FreeBayes v0.9.20 (Garrison and Marth, 2012) and filtered using criteria summarized in Figure 1.

**Figure 1:**
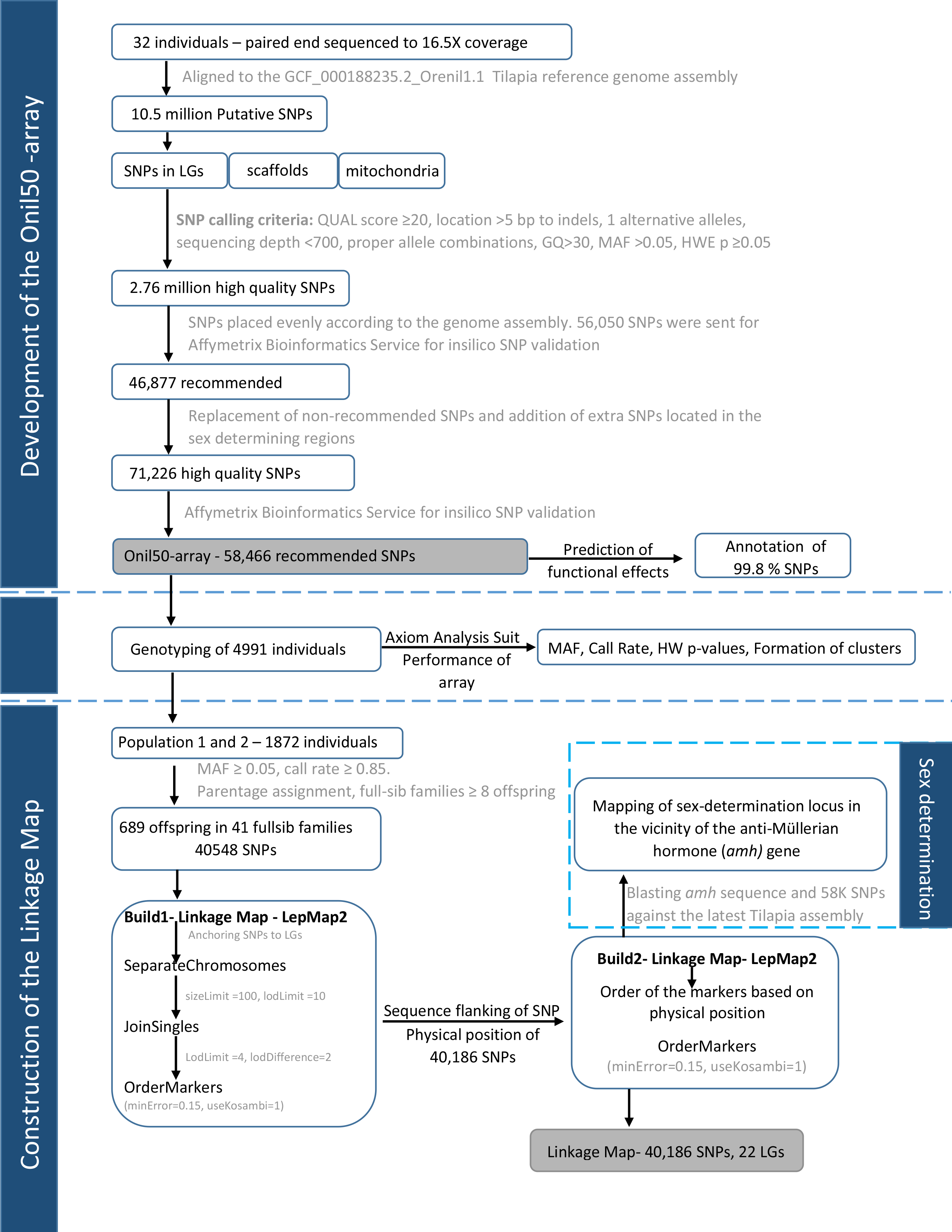
The pipeline for array design, validation and linkage map construction

#### 2.1.2 SNP-filtering

The initial set of putative SNPs was divided into 3 groups including SNPs located on scaffolds assigned to linkage groups, SNPs on unassigned scaffolds, and SNPs detected within the mitochondrial genome. SAMtools v1.2/bcftools (Li et al., 2009) was then used to filter out variants according to a series of criteria. First, as an overall quality filter, SNPs with a QUAL score value (phred) of ≤20 were removed. Then according to the following criteria, a SNP was removed if; (i) located within 5bp to an indel, (ii) had more than one alternative allele, (iii) the sequencing depth exceeded 700 reads, (iv) its alleles were A and T, or C and G, (v) if sample genotype quality was <30, (vi) minor allele frequency (MAF) <0.05, (vii) all samples were heterozygous, (viii) the variant was detected in fewer than 28 of the samples sequenced. Finally, Hardy-Weinberg Equilibrium (HWE) was calculated using PLINK2 (Chang et al., 2015) and SNPs that showed departure from HWE (P<0.05) were removed.

#### 2.1.3 SNP-selection

After filtering, 2.76 million putative high-quality SNPs remained. Based on their relationship to genes and physical distribution, a subset of these were identified for inclusion on the array. SNPEff v 4v1l (Cingolani et al., 2012) was used to identify SNPs with moderate and high effects (including for example non-synonymous variants). From this list of almost 38,000 variants, approximately 10,000 were chosen avoiding SNPs within 10kb of another. A python script was used to fill in gaps and produce a relatively even distribution of SNPs selected at ≈12kb intervals across the 21 linkage groups, and ≈33kb across unmapped scaffolds >50kb in length. The script was designed to fill a distribution gap with a variant falling within a small size selection window with highest minor allele frequency (MAF) being the main criteria. SNPs from the mitochondrial genome were selected manually. The selected subset of SNPs were submitted to *in silico* validation for Affymetrix Bioinformatic Service and based on their recommendation (p-convert value) 58,466 SNPs were chosen to tile on the array. Upon its release, SNP positions were redefined based on the O_niloticus_UMD1 assembly (Conte et al., 2017) using NCBI’s Genome Remapping Service, and SNPEff v 4.3i was rerun to provide updated annotation information.

### 2.2 Genotyping and SNP-performance

Genomic DNA was isolated from ethanol-preserved fin clips collected from 4,991 fish (GST^®^ Tilapia) using Qiagen 96 DNeasy Blood & Tissue Kits according to manufacturer’s instructions (Qiagen, Germany). After quantification and quality checking of DNA, samples were genotyped on the Onil50 array at Center for Integrative Genetics (CIGENE) in Norway. The complete dataset of 4,991 samples was analyzed following the Best Practices Workflow described in Axiom Analysis Suite software (Affymetrix, USA). Four quality parameters were assessed using Axiom Analysis Suit including: MAF, SNP call rates, Hardy Weinberg (HW) p-values, and clustering. With regards to the latter, SNP Polisher classifies SNPs into one of 6 different categories based on cluster profiles with PolyHighRes and NoMinorhom representing the most informative categories. For map construction, only data from SNPs belonging to these categories and displaying a MAF ≥ 0.05 and an overall call rate ≥ 0.85 were used (n= 40,548).

### 2.3 Construction of genetic map

#### 2.3.1 Family structure

Genotypes from a subset of 1872 samples was used for map construction. Population 1 (n=1124) were individuals collected following the branching of the 20th generation, and were factorially crossed against each other after 2 generations. The experimental design for Population 1 is described in Joshi et al. (2018) and was primarily intended to partition the non-additive genetic effects in this population. Population 2 (n=748) were obtained from the 24th and 25th generations of GST^®^.

Parentage assignment was done using an exclusion method which eliminates animals from a list of potential parents when there are opposing homozygotes between parents and offspring (Hayes, 2011). We permitted a maximum of 100 conflicts between parents and offspring, representing approximately 0.24% of all genotypes. A total of 689 offspring were divided among 41 full-sib families containing ≥ 8 offspring (mean offspring per family, μ=16.81). Population 1 (468 offspring + 19 parents) had 34 full-sib families (μ = 13.77 ± 5.5) and Population 2 (221 offspring + 14 parents) had 7 full-sub families (μ = 31.57 ± 7.23). The structure of Population 1 and 2 is shown in Supplementary Table 1 and 2.

#### 2.3.2 Linkage map construction

Phenotypic sex were known for a subset of families (221 offspring + 33 parents) and was coded as 12 for males and 11 for females and included in the genotype file before running Lepmap2 (Rastas et al., 2013) for linkage map construction. Lepmap2 uses information from full-sibs to assign SNP markers to linkage groups (LGs), and applies standard hidden Markov model (HMM) to compute the likelihood of the marker order within each LGs. First, the SNPs were used to construct the preliminary linkage map (Build 1), which was used to anchor, order and orient the scaffolds in the O_niloticus_UMD1 assembly and upgrading this assembly to O_niloticus_UMD_NMBU (Conte et al., 2018). Eventually, the final physical integrated genetic linkage map (Build 2) was constructed from the order of the markers based on the physical position of the SNPs in O_niloticus_UMD_NMBU assembly.

#### 2.3.3 Build 1: To anchor SNPs to different LGs

SeparateChromosomes (a module in Lepmap2) was run testing lodLimits from 1 - 50 and a sizeLimit = 100; a lodLimit of 10 resulted in 22 LGs with lowest number of markers not assigned to any LG. JoinSingles was run to assign single markers to LG groups and tested with lodLimits from 1 - 15 and lodDifference = 2; a lodLimit of 4 was selected as this joined the highest number of single markers. OrderMarkers was used to order the markers within each LG. Each LG was ordered separately and replicated 5 times with commands: numThreads=10 polishWindow=30 filterWindow=10 useKosambi=1 minError=0.15, and the order with highest likelihood was selected as the best order. For sex averaged map OrderMarkers was run similarly by adding sexAverage=1.

#### 2.3.4 Build 2: Integrated linkage map based on the order of the SNPs in the new assembly

Sequence flanking each SNP was used to find the physical position of 40,186 SNPs in the O_niloticus_UMD_NMBU assembly. Physical position information was used to adjust the order of the SNPs within respective linkage groups and Lepmap2 was rerun to produce the final linkage map.

## 3 Results

### 3.1 SNP selection and array development

The sequencing of 32 tilapia generated 528 million reads representing an average of 16.5x coverage per individual. On average 98% of reads were mapped to the Orenil1.1 assembly yielding 12.7 million variants of which 10.5 million were putative SNPs. After performing multiple steps of filtering based on a markers behavior and amenability to Axiom technology, a subset of 2.76 million SNPs were retained and further filtered to produce a final set of 58,466 SNPs for which assays were designed and printed in the 0nil50 array.

The assignment of SNPs to linkage groups and unmapped scaffolds in Orenil 1.1 (used for SNP selection) and O_niloticus_UMD1 with 99.8% of the SNPs were successfully re-mapped to the new assembly (Table 1). Remapping revealed an increase in the number of SNPs mapping to linkage groups and a corresponding decrease in the number of SNPs on unmapped scaffolds. The average variant density per linkage group on the Orenil1.1 assembly is 12,5kb. However, since the O_niloticus_UMD1 assembly includes an additional 87 Mb assigned to LGs the average distance between the variants increased to 15,5kb in this assembly. The most significant difference is a 2.4 fold increase in the physical map size for LG 3 which produced a 2.3 fold increase in the number of SNPs assigned to this linkage group.

**Table 1:**
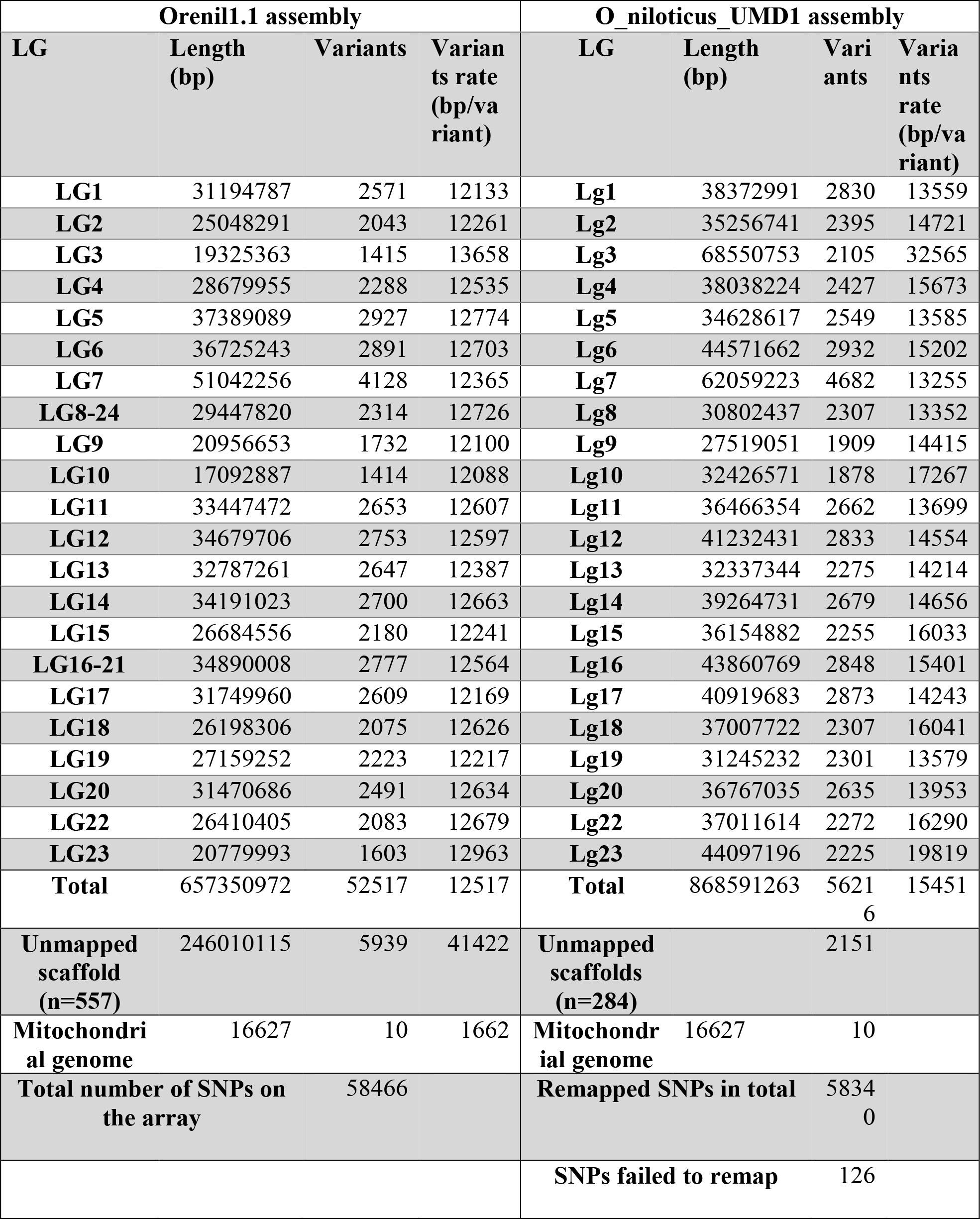
Sequence similarity based assignment of SNPs contained on the 0nil50-array to Orenill.1 and O_niloticus_UMD1 genome assemblies.

### 3.2 Performance and validation of the SNPs in the array

Performance of the SNPs on the array was further explored with genotyping of 4,858 additional tilapia samples with call rates exceeding 80%. Axiom Analysis Suite was used to categorize the SNPs into classes reflecting their cluster profiles. Over 74% of the SNPs were classified as PolyHighResolution. More detailed information about the sample and SNP statistics are shown in Figure 2.

**Figure 2:**
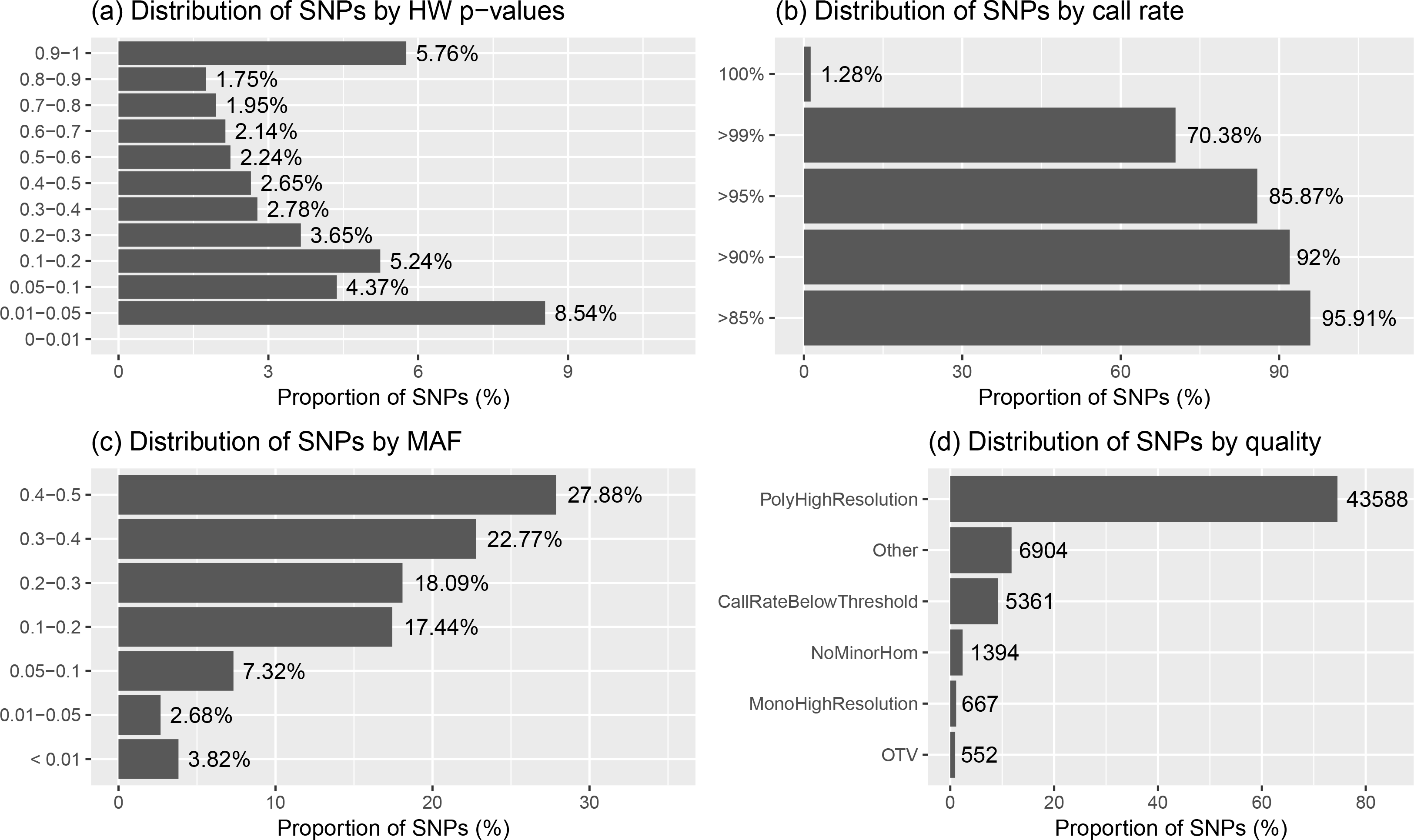
Summary of the SNP metrics based on HW p-values, call rate, MAF and the types of clusters

Running snpEffv4.3i (Cingolani et al., 2012) to predict functional effects of the 58,340 remapped SNPs from the Onil50 array resulted, in most cases, in multiple annotations per variant. The effects with the highest putative impact are included for summary in Table 2. The majority of the SNPs are intronic or intergenic variants, while about 15% of are nonsynonymous mutations. These variants can have direct effect on a trait of interest and are a direct result of the SNP selection process which specifically targeted variants with a potential functional effect.

**Table 2:** Summary of annotation for SNPs in the Onil50-array

| SNP categories | Count | Percent |
| --- | --- | --- |
| **Total number of SNPs in the array** | 58,446 |  |
| **Annotation Possible** | 58,340 | 99.82 |
| **Annotation results** |  |  |
| Nonsense-mediated decay (NMD) | 19 | 0.03 |
| Loss of function (LOF) | 114 | 0.20 |
| Intergenic region | 12,156 | 20.80 |
| Intragenic variant | 126 | 0.22 |
| Intron variant | 21,581 | 36.92 |
| **Non-synonymous variant** |  |  |
| Missense variant | 8,765 | 15.00 |
| Missense variant & splice region variant | 263 | 0.45 |
| Stop gained | 16 | 0.03 |
| Stop lost | 13 | 0.02 |
| Synonymous variant | 1,142 | 1.95 |
| Non coding transcript exon variant | 27 | 0.05 |
| Splice acceptor variant & intron variant | 9 | 0.02 |
| Splice donor variant & intron variant | 13 | 0.02 |
| Splice region variant | 8 | 0.01 |
| Splice region variant & intron variant | 163 | 0.28 |
| Splice region variant & non coding transcript exon variant | 7 | 0.01 |
| Splice region variant & synonymous variant | 28 | 0.05 |
| Upstream gene variant | 9,231 | 15.79 |
| 3 prime UTR variant | 1,533 | 2.62 |
| 5 prime UTR premature start codon gain variant | 21 | 0.04 |
| 5 prime UTR variant | 459 | 0.79 |
| Downstream gene variant | 2,646 | 4.53 |

### 3.3 Linkage Map

A total of 40,549 SNPs were retained following quality filtering, and 99.78% of these (n = 40,467) were ordered within the 22 linkage groups corresponding to the karyotype of Nile tilapia (Supplementary Figure 1). Since, Build 1 linkage map is an intermediate step for the extension of the 0_niloticus_UMD1 to the 0_niloticus_UMD_NMBU genome assembly (Conte et al., 2018), and this not the aim of this paper, we give only a brief summary of the results. The genetic and physical maps were generally in good agreement with a correlation of ≥ 0.96 between the reference genome position and the genetic map position of the SNPs (Supplementary Figure 1). This high correlation with the physical map demonstrates that the genetic map is of high quality and is highly accurate.

A total of 40,186 SNPs mapped to 22 linkage groups in Build 2 linkage map. The consensus (sex-averaged) map adds up to 1469.69 cM, with individual linkage group lengths ranging from 56.04 cM (LG19) to 96.68 cM (LG07) (Table 3). The average genetic distance across the LGs was 66.8 cM. The number of markers per LG varied from 1349 to 3391, with an average of 1827 markers per LG (Table 3). As a consequence of the SNP selection, which sought to position a SNP every 12kb, the number of markers was mostly proportional to the size of the LG (Figure 4). A notable exception is LG03 where the inclusion of previously unassigned scaffolds has trippled the physical size without a corresponding tripling of SNP numbers. The SNP density (cM/locus) varied across the genome, which can be seen in Figure 3 and Supplementary Figures 2–4.

**Figure 3:**
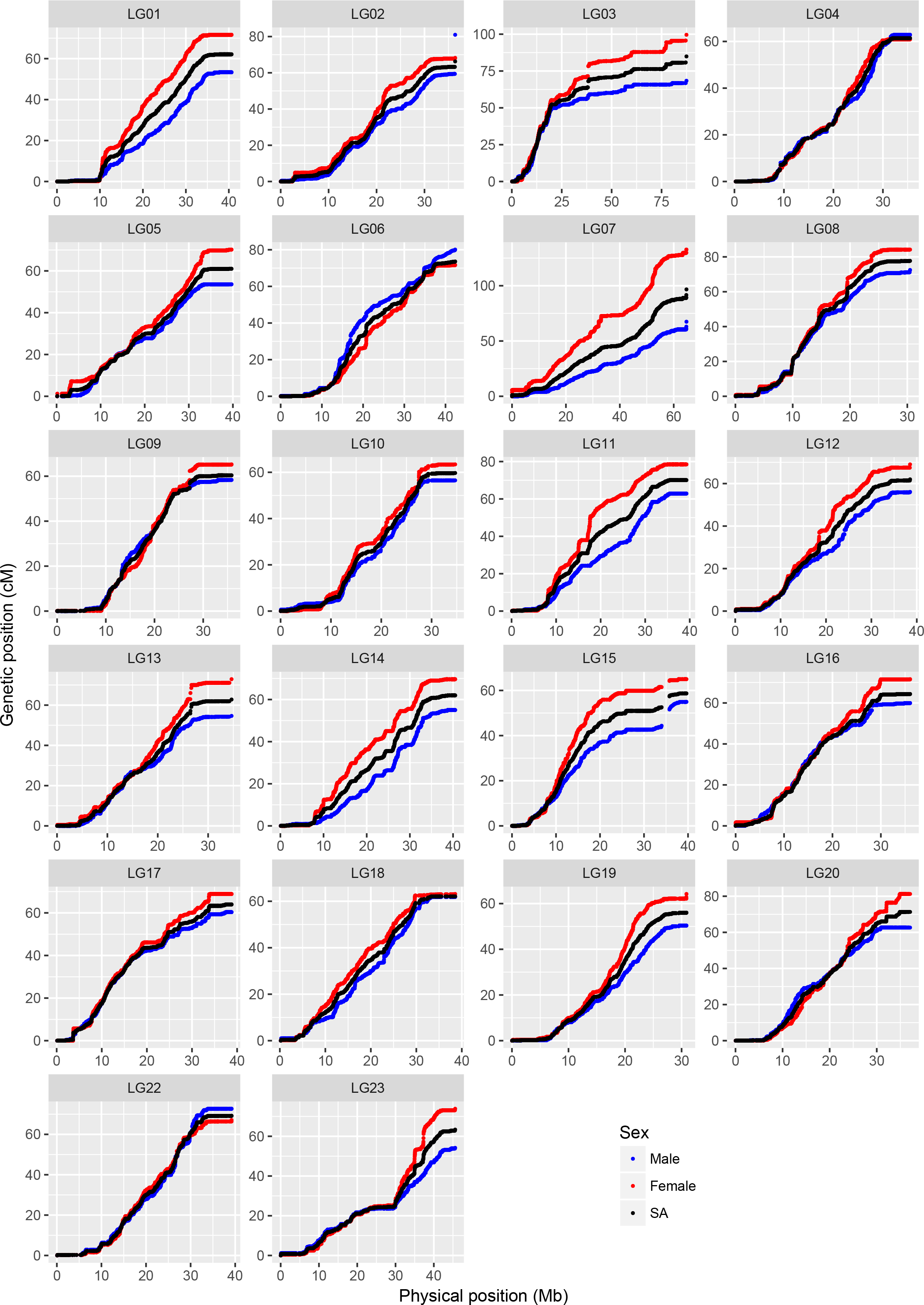
Comparison of map positions between genetic and physical maps for different LGs in Build 2. The y-axis gives the linkage map positions, and the x-axis gives the physical positions. Linkage groups and the physical positions are based on O_niloticus_UMD_NMBU Assembly. The maps are color-coded: red for female specific, blue for male specific and black for sex-averaged linkage maps.

**Figure 4:**
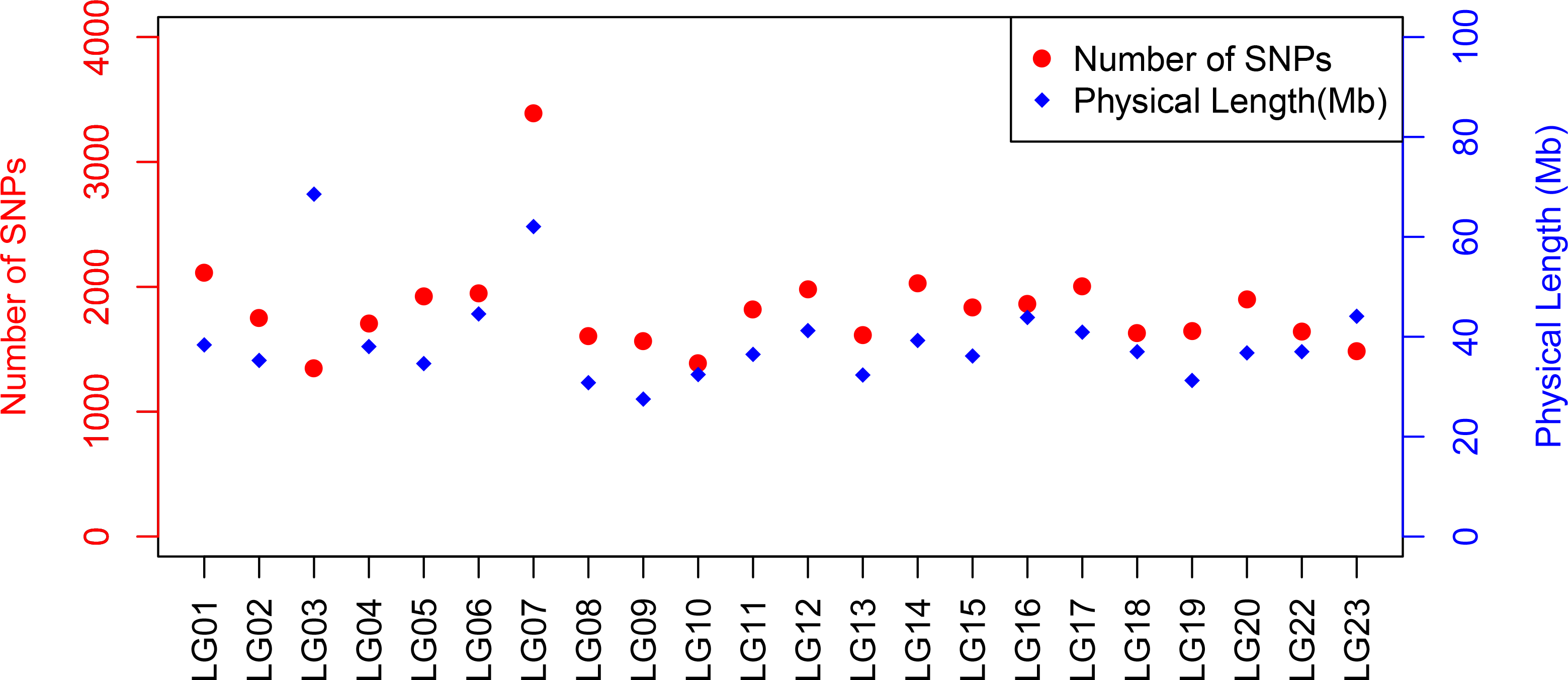
Plot illustrating the number of SNPs and physical length of linkage group based on O_niloticus_UMD1 Assembly and Build 2 linkage map.

**Table 3:**
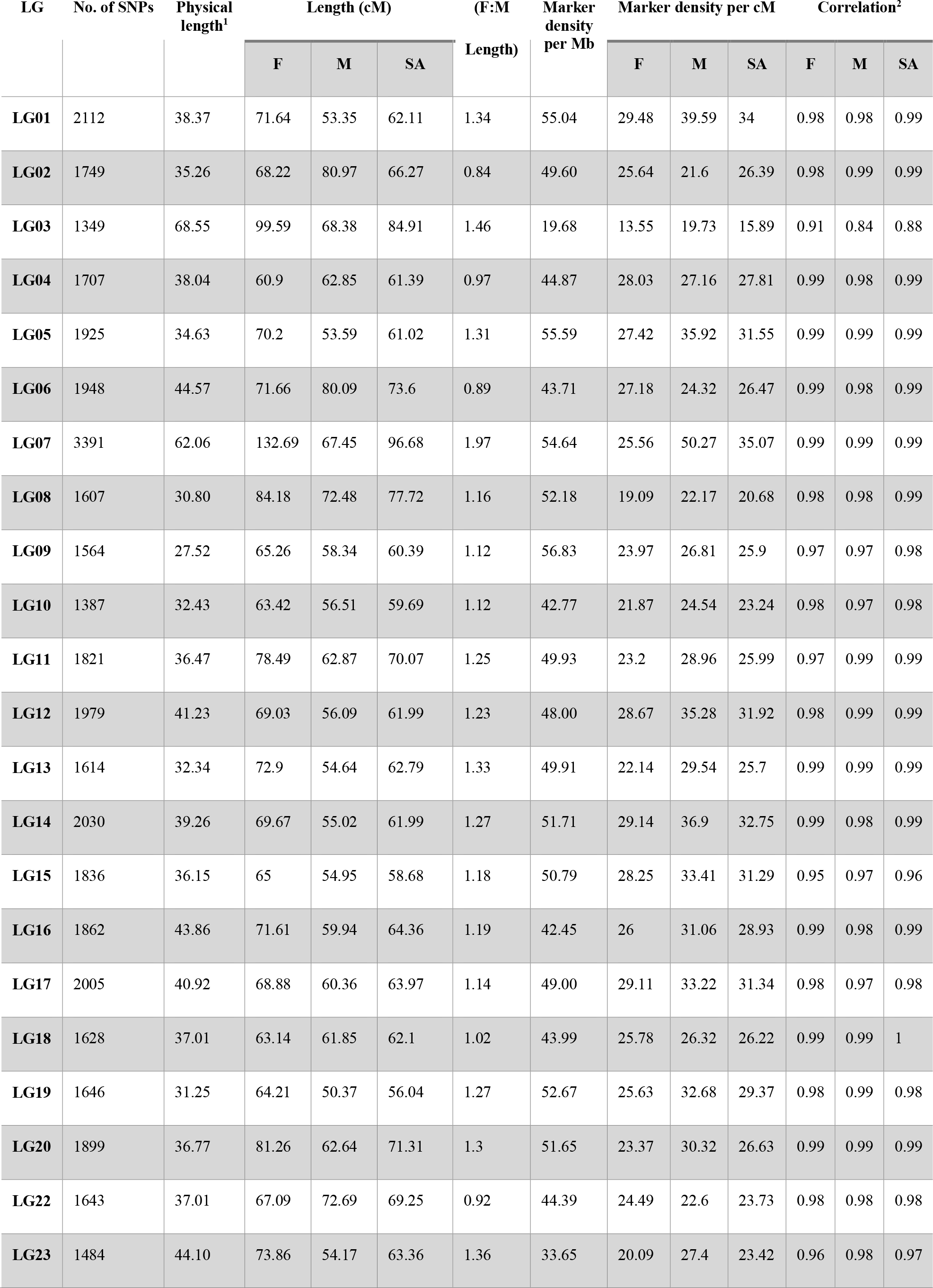
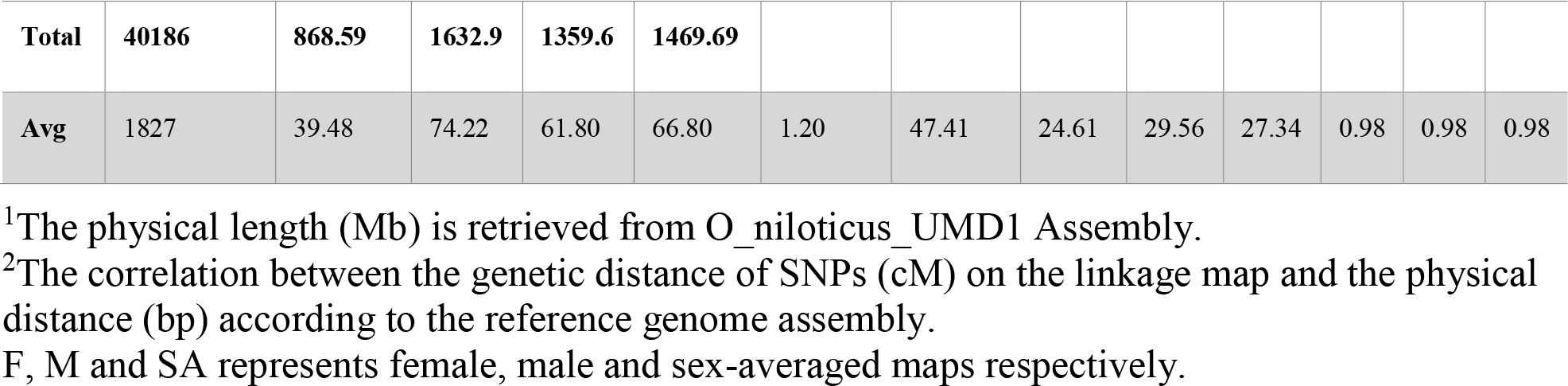
Marker numbers, length, density and correlations for male, female and sex-averaged Build 2 linkage map

| LG | No. of SNPs | Physical length<sup>1</sup> | Length (cM) |  |  | (F:M Length) | Marker density per Mb | Marker density per cM |  |  | Correlation<sup>2</sup> |  |  |
| --- | --- | --- | --- | --- | --- | --- | --- | --- | --- | --- | --- | --- | --- |
|  |  |  | F | M | SA |  |  | F | M | SA | F | M | SA |
| LG01 | 2112 | 38.37 | 71.64 | 53.35 | 62.11 | 1.34 | 55.04 | 29.48 | 39.59 | 34 | 0.98 | 0.98 | 0.99 |
| LG02 | 1749 | 35.26 | 68.22 | 80.97 | 66.27 | 0.84 | 49.60 | 25.64 | 21.6 | 26.39 | 0.98 | 0.99 | 0.99 |
| LG03 | 1349 | 68.55 | 99.59 | 68.38 | 84.91 | 1.46 | 19.68 | 13.55 | 19.73 | 15.89 | 0.91 | 0.84 | 0.88 |
| LG04 | 1707 | 38.04 | 60.9 | 62.85 | 61.39 | 0.97 | 44.87 | 28.03 | 27.16 | 27.81 | 0.99 | 0.98 | 0.99 |
| LG05 | 1925 | 34.63 | 70.2 | 53.59 | 61.02 | 1.31 | 55.59 | 27.42 | 35.92 | 31.55 | 0.99 | 0.99 | 0.99 |
| LG06 | 1948 | 44.57 | 71.66 | 80.09 | 73.6 | 0.89 | 43.71 | 27.18 | 24.32 | 26.47 | 0.99 | 0.98 | 0.99 |
| LG07 | 3391 | 62.06 | 132.69 | 67.45 | 96.68 | 1.97 | 54.64 | 25.56 | 50.27 | 35.07 | 0.99 | 0.99 | 0.99 |
| LG08 | 1607 | 30.80 | 84.18 | 72.48 | 77.72 | 1.16 | 52.18 | 19.09 | 22.17 | 20.68 | 0.98 | 0.98 | 0.99 |
| LG09 | 1564 | 27.52 | 65.26 | 58.34 | 60.39 | 1.12 | 56.83 | 23.97 | 26.81 | 25.9 | 0.97 | 0.97 | 0.98 |
| LG10 | 1387 | 32.43 | 63.42 | 56.51 | 59.69 | 1.12 | 42.77 | 21.87 | 24.54 | 23.24 | 0.98 | 0.97 | 0.98 |
| LG11 | 1821 | 36.47 | 78.49 | 62.87 | 70.07 | 1.25 | 49.93 | 23.2 | 28.96 | 25.99 | 0.97 | 0.99 | 0.99 |
| LG12 | 1979 | 41.23 | 69.03 | 56.09 | 61.99 | 1.23 | 48.00 | 28.67 | 35.28 | 31.92 | 0.98 | 0.99 | 0.99 |
| LG13 | 1614 | 32.34 | 72.9 | 54.64 | 62.79 | 1.33 | 49.91 | 22.14 | 29.54 | 25.7 | 0.99 | 0.99 | 0.99 |
| LG14 | 2030 | 39.26 | 69.67 | 55.02 | 61.99 | 1.27 | 51.71 | 29.14 | 36.9 | 32.75 | 0.99 | 0.98 | 0.99 |
| LG15 | 1836 | 36.15 | 65 | 54.95 | 58.68 | 1.18 | 50.79 | 28.25 | 33.41 | 31.29 | 0.95 | 0.97 | 0.96 |
| LG16 | 1862 | 43.86 | 71.61 | 59.94 | 64.36 | 1.19 | 42.45 | 26 | 31.06 | 28.93 | 0.99 | 0.98 | 0.99 |
| LG17 | 2005 | 40.92 | 68.88 | 60.36 | 63.97 | 1.14 | 49.00 | 29.11 | 33.22 | 31.34 | 0.98 | 0.97 | 0.98 |
| LG18 | 1628 | 37.01 | 63.14 | 61.85 | 62.1 | 1.02 | 43.99 | 25.78 | 26.32 | 26.22 | 0.99 | 0.99 | 1 |
| LG19 | 1646 | 31.25 | 64.21 | 50.37 | 56.04 | 1.27 | 52.67 | 25.63 | 32.68 | 29.37 | 0.98 | 0.99 | 0.98 |
| LG20 | 1899 | 36.77 | 81.26 | 62.64 | 71.31 | 1.3 | 51.65 | 23.37 | 30.32 | 26.63 | 0.99 | 0.99 | 0.99 |
| LG22 | 1643 | 37.01 | 67.09 | 72.69 | 69.25 | 0.92 | 44.39 | 24.49 | 22.6 | 23.73 | 0.98 | 0.98 | 0.98 |
| LG23 | 1484 | 44.10 | 73.86 | 54.17 | 63.36 | 1.36 | 33.65 | 20.09 | 27.4 | 23.42 | 0.96 | 0.98 | 0.97 |
| **Total** | **40186** | **868.59** | **1632.9** | **1359.6** | **1469.69** |  |  |  |  |  |  |  |  |
| **Avg** | 1827 | 39.48 | 74.22 | 61.80 | 66.80 | 1.20 | 47.41 | 24.61 | 29.56 | 27.34 | 0.98 | 0.98 | 0.98 |

<sup>1</sup>The physical length (Mb) is retrieved from O\_niloticus\_UMD1 Assembly.
<sup>2</sup>The correlation between the genetic distance of SNPs (cM) on the linkage map and the physical distance (bp) according to the reference genome assembly.
F, M and SA represents female, male and sex-averaged maps respectively.

In this study, paternal and maternal informative markers were used to construct specific male and female maps. (Table 3). Around one-third of the linkage groups showed a different recombination rate between sexes, with male and female map lengths differing by a factor of 1.2 (1359.6cM and 1632.9cM respectively). Generally female maps were found to be larger, with the exception of LG02, LG06 and LG22. Sigmoidal pattern of recombination, with no recombination at both ends of the LGs, was seen in almost all linkage groups (Figure 3).

## 4 Discussion

### 4.1 High-density linkage map for Tilapia

Existing linkage maps for Nile tilapia contain relatively few markers unevenly distributed across linkage groups (Supplementary Table 3). As a consequence, regions in the genome have poor SNP coverage. By stringently selecting SNPs with an even physical distribution in the genome the linkage map presented includes 10 times more SNPs and fewer gaps.

Ferreira et al. (2010) categorized the karyotypes of *O. niloticus* into 3 meta-submetacentric and 19 subtelo-acrocentric chromosomes. The steepness of the curve in Figure 3 shows the recombination level, with flat lines representing little or no recombination, which may suggest the possible location of the centromeres. These notable features, i.e. the wide recombination deserts (areas with no recombination), are seen in the initial and/or end regions of most of the linkage groups, generally up to 5 Mb and sometimes up to 10 Mb (e.g. LG09 and LG10), revealing the presence of mainly subtelo-acrocentric linkage groups. Because of these recombination deserts, most of the linkage groups, irrespective of the sexes, showed sigmoidal pattern, which is unusual when compared to other fish species. In channel catfish (Li et al., 2014), salmon (Tsai et al., 2015), Asian seabass (Wang et al., 2015) and stickleback (Roesti et al., 2013) the recombination rates were generally elevated towards the end of the linkage groups. The possible explanation might be that the GST^®^ strain used in this study is derived from the GIFT strain, formed from crossing among four wild and four cultured Asian strains (Eknath et al., 1993), which might have given us the unique recombination pattern.

Tilapia have been shown to have a sex-specific pattern of recombination with the female map generally being larger than the male map (Lee et al. 2004). The genetic basis for the differences in the recombination in different sexes has still not been found, but Li et al. (2014) has listed three major hypotheses. First, the selection perspective hypothesis (Gruhn et al., 2013; Lenormand and Dutheil, 2005), proposes that the selection pressure is higher in male gametes compared to female gametes during the haploid life stage and this male-specific selection leads to decrease in the male recombination rate to maintain the beneficial haplotypes. Secondly, the compensation hypothesis (Coop and Przeworski, 2007), proposes that the recombination rate is higher in females compared to males to compensate for the less stringent checkpoint for the achiasmatic chromosomes. Thirdly, the recombination pathway hypothesis (Gruhn et al., 2013), suggests that the chromatin differences established prior to the onset of the recombination pathway causes the differences in the recombination between the two sexes.

LG23 showed a unique recombination pattern, a flat line of around 5 Mb, in the centre of the linkage group, for which there also is a sex difference in recombination rate. In *O. niloticus,* major XY sex determining regions have earlier been mapped to LG1 (Palaiokostas et al., 2013) and LG23 (Eshel et al., 2011, 2012; Karayücel et al., 2004; Shirak et al., 2006). Further, tandem duplication of the variants of the gene anti-Müllerian hormone (*amh*) in LG23 has been identified as as the male sex determinant in Tilapia (Li et al., 2015). These variants of *amh* gene have been mapped to around 35.4 Mb region of tilapia genome (discussed below in section 4.3), which is the same region where the unique recombination pattern is seen, suggesting limited recombination around the sex determining genes in *O. niloticus.* Further, LG23 was formed by the fusion of two linkage groups during the evolution of cichlids (Liu et al., 2013), which might be another reason for this unique recombination pattern.

The fusion of the linkage groups during the evolutionary process also has an effect on the size of the linkage groups, as it is believed that the ancestors of cichlids had 24 chromosome pairs, which eventually became 22 pairs (Majumdar and McAndrew, 1986). The physical map and the cytogenic studies indicate LG03 and LG07 consequently became the two largest linkage groups (Conte et al., 2017; Ferreira et al., 2010; Liu et al., 2013; Poletto et al., 2010), which is also supported by our genetic map.

### 4.2 Array content and performance

SNP performance was validated by genotyping around 5000 individuals from different generations of the GST^®^ strain of tilapia. Around 75% of the SNPs on the array perform well generating three highly differentiated allelotype clusters (i.e. polyhighresolution). Around 9% of the SNPs were found to depart from HWE (p < 0.01), but it has to be noted that the population genotyped for the validation is the commercial strain that has undergone up to 25 generations of selection. Hence, these departures might be important as they could represent regions under selection and the outcome of assortative mating. Whereas the extreme departures might suggest lethal recessive mutations and/or recent mutations or copy number variants.

For future revisions, the array could be improved by increasing the SNP density in highly recombinant regions of specific linkage groups like including LG03 and LG23. The use of genetic distance rather than the physical distance to select the SNPs is probably the best option for equidistant SNP distribution across the genome.

### 4.3 Sex locus mapped in the vicinity of *amh* gene

Sex determination is one of the important aspect in commercial tilapia production, as males are found to grow faster than females and unisex production is a main method to avoid propagation in production ponds or cages. Sex determination in fish is more complicated than mammals as it tends to be co-dependent on both genetic and environmental factors (Baroiller et al., 2009; Ezaz et al., 2006). Two main sex determination system exists: XY and ZW, and they are both present in fish species. It has also been seen that closely related fish species, even in same genus, have different sex determination systems. For example, Blue tilapia, *Oreochromis aureus,* has the ZW system of sex determination (Campos-Ramos et al., 2001), where males are homogametic (ZZ) and females are heterogametic (ZW), so the ovum determines the sex of the offspring. Whereas, Nile tilapia (*O. niloticus*) and Mozambique tilapia (*O. mossambicus*) have the XY system of sex determination, where the males are heterogametic (XY) and females are homogametic (XX), so the sperm determines the sex of the offspring (Campos-Ramos et al., 2003; Mair et al., 1991).

In our study, the sex locus of tilapias was coded using the XY system and mapped to LG23 (Table 4) as reported previously in several studies (Eshel et al., 2011, 2012; Karayücel et al., 2004; Shirak et al., 2006). The most likely position of sex locus (pos. 34.5Mb/40.53 cM on LG23) maps close to the anti-Müllerian hormone (*amh*) gene, previously characterized as sex determining gene in Nile tilapia (Li et al., 2015).

**Table 4:** Mapping of sex-determination locus in the vicinity of the anti-Mullerian hormone *(amh)* gene. SNP AX-164998274 (marked as *) mapped to the same genetic position as the Phenotypic sex of the individuals in the Build 1 linkage map.

| SNPs/gene | LG | Position (bp) | Male | Female | Average |
| --- | --- | --- | --- | --- | --- |
| AX-165032341 | LG23 | 34305951 | 35.03 | 44.83 | 39.97 |
| AX-164990538 | LG23 | 34306186 | 35.03 | 44.83 | 39.97 |
| AX-165017655 | LG23 | 34319855 | 35.03 | 44.83 | 39.97 |
| AX-165032969 | LG23 | 34336514 | 35.03 | 44.83 | 39.97 |
| AX-165012489 | LG23 | 34351488 | 35.03 | 44.83 | 39.97 |
| AX-164995826 | LG23 | 34367182 | 35.24 | 44.83 | 40.00 |
| AX-165001648 | LG23 | 34380102 | 35.45 | 44.83 | 40.09 |
| AX-165030187 | LG23 | 34380282 | 35.45 | 44.85 | 40.12 |
| AX-164992183 | LG23 | 34398468 | 35.45 | 44.88 | 40.13 |
| AX-165006758 | LG23 | 34424845 | 35.45 | 44.95 | 40.15 |
| AX-164986178 | LG23 | 34437472 | 35.45 | 45.02 | 40.20 |
| AX-165024637 | LG23 | 34451454 | 35.45 | 45.61 | 40.53 |
| AX-165013086 | LG23 | 34465412 | 35.45 | 45.61 | 40.53 |
| AX-164998274* | LG23 | 34496900 | 35.45 | 45.61 | 40.53 |
| **amh_delta-y** | **LG23** | **34491516-34499598** |  |  |  |
| **amhy** | **LG23** | **34491516-34503495** |  |  |  |
| **amh** | **LG23** | **34491516-34509687** |  |  |  |
| AX-164990628 | LG23 | 34510978 | 35.45 | 45.61 | 40.53 |
| AX-165031999 | LG23 | 34511701 | 35.46 | 45.61 | 40.54 |
| AX-165013176 | LG23 | 34525091 | 35.46 | 45.61 | 40.54 |
| AX-165010851 | LG23 | 34576386 | 35.46 | 45.61 | 40.54 |
| AX-164993854 | LG23 | 34585587 | 35.46 | 45.61 | 40.54 |
| AX-164989444 | LG23 | 34598712 | 35.46 | 45.61 | 40.54 |

### 4.4 Implications in commercial tilapia industry

Tilapia is a commercially important aquaculture species, surpassing salmon in terms of production, with more than 3.9 million tons of fish and fillets being traded in 2015 (FAO, 2017) and more than 20 breeding programs (Neira, 2010). The present SNP array and linkage map has the potential to greatly improve the genetic gain for this economic important species, and help surpass the difficulties of efficient selection for the invasive traits, the traits which can’t be measured directly on the candidate broodstock fish, but are only measured on the sibs of the candidates e.g. disease resistance, fillet yield, etc. These tools may also be useful to bridge the genotype-phenotype gap in Nile tilapia, which has been pursued for a long time (Gjøen, 2004).

A major capability of these resources will be to find economic important QTLs or chromosome regions affecting economically important traits like disease resistance, fillet traits or feed efficiency. In order to fine map these QTLs, it is essential to have a high-resolution linkage map. The dense linkage map can also be integrated with physical maps to position and orient scaffolds along linkage groups, thereby producing genome assemblies of higher quality.

Another important implication will be to facilitate the shift from traditional breeding strategies to genomic selection in Nile tilapia. The breeding goals in Tilapia will in the future include many invasive traits. Genomic selection will significantly help us to overcome these challenges, increasing the profitability and the genetic gain (Hosoya et al., 2017; Houston, 2017; Meuwissen et al., 2001; Nielsen et al., 2009; Sonesson and Meuwissen, 2009; Vela-Avitúa et al., 2015). Finally, this will also help to separate the additive and non-additive genetic effects more accurately, thereby increasing both the accuracy of the selection and the possibility to utilize non-additive genetic effects (Varona et al., 2018).

Another obvious use of the SNP-array will be in the parentage assignment. The drawback of the conventional breeding designs in Tilapia using PIT tags is the confounding of the full-sib family effects (due to communal rearing of full-sibs) and maternal environmental effects (due to mouth brooding), making it difficult to detangle the various variance components accurately (Joshi et al., 2018), which ultimately decreases the accuracy of the selection.

## 5 Conclusion

We present the first SNP-array, the Onil50-array, containing ca 58,000 SNPs for Nile Tilapia, which was validated in 5000 individuals. Further, we constructed a high density integrated genetic and physical linkage map, with linkage groups showing sex-differentiated sigmoidal recombination patterns. These new resources has the potential to greatly influence and improve the genetic gain when applying genomic selection and surpass the difficulties of efficient selection for invasive traits in tilapia.

## 8 Author Contributions

HG, AA and MK conceived and designed the study. AA coordinated biological sampling. MK and MA were responsible for array design and MA performed lab work and initial analysis of results. RJ constructed the linkage map, while SL integrated the genetic and physical maps. RJ and MA prepared the draft manuscript which was reviewed and edited by HG, MK, AA and SL. All authors read and approved the manuscript.

## 10 Acknowledgments

We would like to acknowledge Anders Skaarud from Genomar Genetics AS. We would like to thank Harald Grove, Torfinn Nome and Tim Knutsen for their valuable advice and help with bioinformatics analyses of sequence and SNP data. The SNP array was developed in cooperation with Affymetrix, Inc., and we particularly thank the following Affymetrix personnel for their direct contribution: Lakshmi Radhakrishnan and Alessandro Davassi. Similarly, we also acknowledge Solomon Boison and Luqman Aslam for their interaction during linkage map construction.

## 11 Data Availability

The whole genome sequence data used for SNP detection was generated using broodstock from a breeding population and is commercially sensitive. Similarly, the genotypes used for map construction are from commercial family material. This information may be made available to noncompetitive interests under conditions specified in a Data Transfer Agreement. Requests to access these datasets should be directed to Alejandro Tola Alvarez.

The assemblies used in this study can be found in NCBI using the following accessions: Orenil1.1= GCF_000188235.2, O_niloticus_UMD1 = MKQE00000000 and O_niloticus_UMD_NMBU = MKQE02000000. Linkage map generated from this study can be found in the Figshare: https://figshare.com/s/8427b97cf6e623173232

## 13 Supplementary Material

Supplementary file containing Supplementary Tables and Figures follows this manuscript.

**Supplementary Table 1:** Observations in each factorial mating in Population 1. 11 different sires (M1 to M11) are mated with 8 different dams (F1 to F8) in factorial manner. Only those full-sib families ≥ 8 offspring are shown.

|  | **F1** | **F2** | **F3** | **F4** | **F5** | **F6** | **F7** | **F8** | **Total** |
| --- | --- | --- | --- | --- | --- | --- | --- | --- | --- |
| **M1** | 22 | 13 | 10 | - | 8 | - | - | - | **53** |
| **M2** | - | - | - | 10 | - | 21 | 10 | 24 | **65** |
| **M3** | - | - | - | 12 | - | 23 | 11 | 16 | **62** |
| **M4** | - | - | - | 13 | - | 9 | - | - | **22** |
| **M5** | 11 | 24 | 9 | - | 8 | - | - | - | **52** |
| **M6** | - | - | - | - | - | 18 | 8 | 13 | **39** |
| **M7** | 19 | 12 | - | - | - | - | - | - | **31** |
| **M8** | - | - | - | 14 | - | 9 | 12 | 28 | **63** |
| **M9** | - | 14 | - | - | 10 | - | - | - | **24** |
| **M10** | 16 | 14 | 8 | - | 8 | - | - | - | **46** |
| **M11** | - | - | - | 11 | - | - | - | - | **11** |
| **Total** | **68** | **77** | **27** | **60** | **34** | **80** | **41** | **81** | **468** |

**Supplementary Table 2:** Observations in different full-sib families in the Population 2

| **Dam** | **F9** | **F10** | **F11** | **F12** | **F13** | **F14** | **F15** | **Total** |
| --- | --- | --- | --- | --- | --- | --- | --- | --- |
| **Sire** | **M12** | **M13** | **M14** | **M15** | **M16** | **M17** | **M18** |  |
| **No. of offspring** | 22 | 24 | 26 | 36 | 37 | 37 | 39 | **221** |

**Supplementary Table 3:**
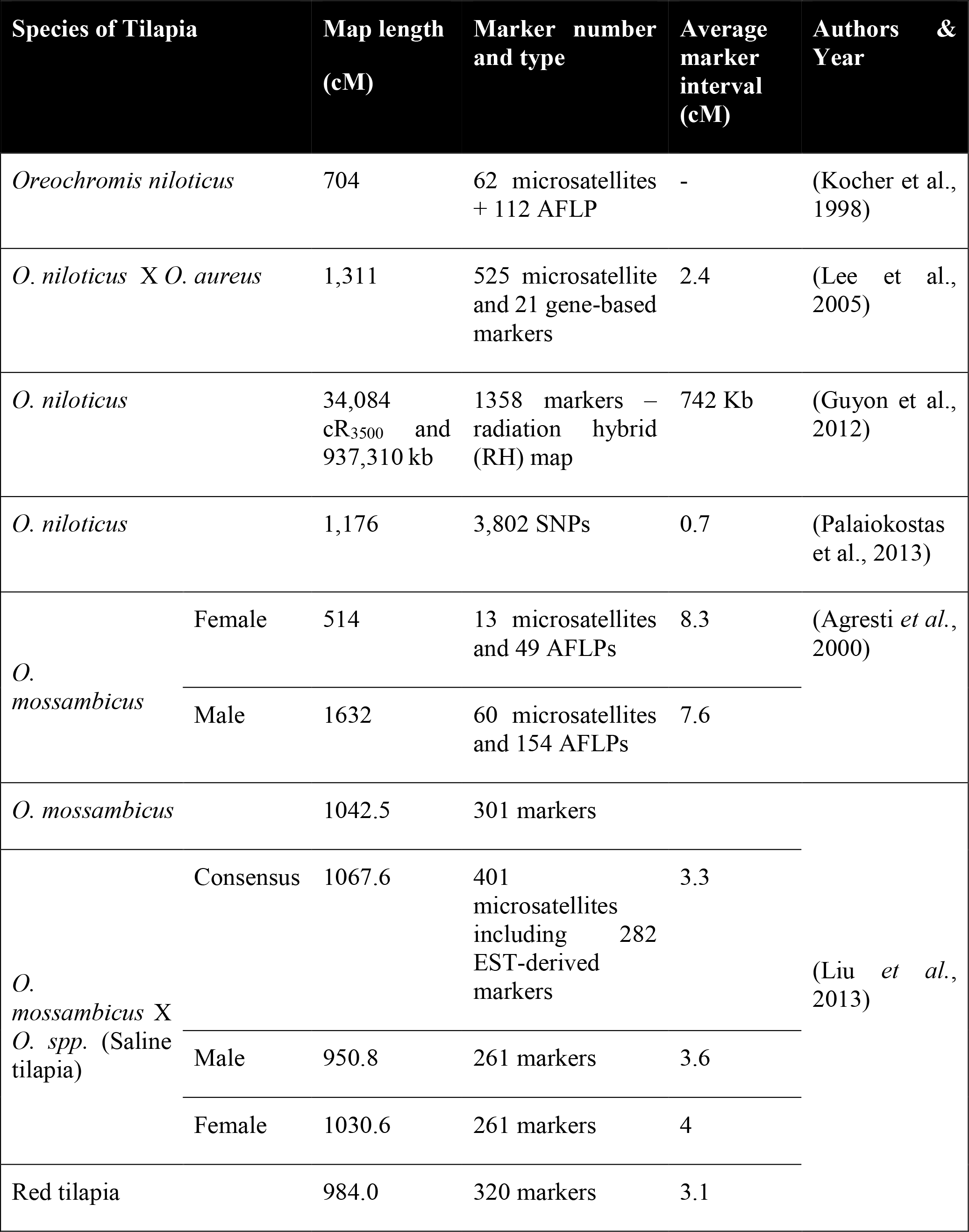
Published linkage maps for Tilapia species

**Supplementary Figure 1:**
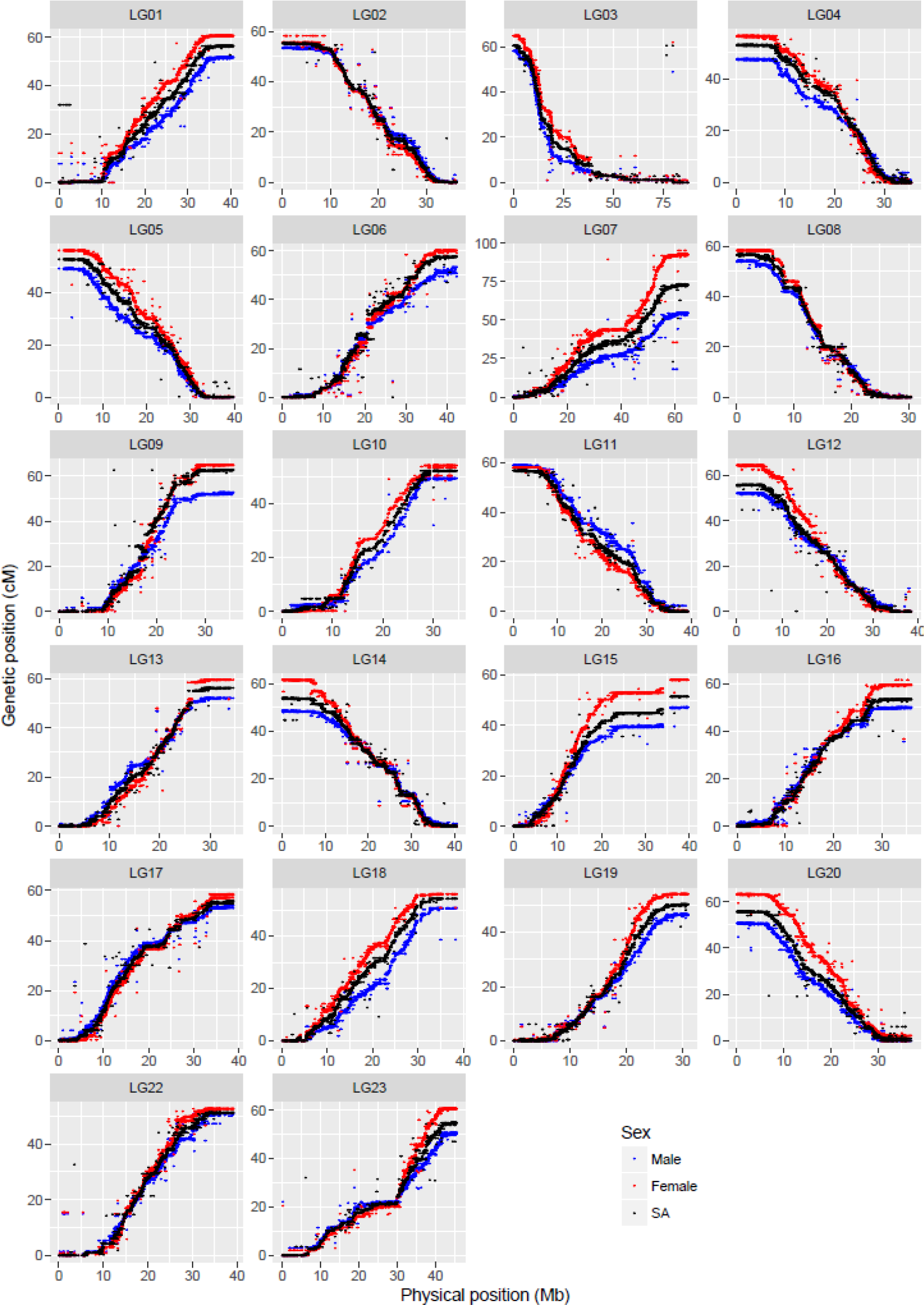
Comparison of map positions between genetic and physical maps for different LGs in Build1 linkage map. The y-axis gives the linkage map positions, and the x-axis gives the physical positions. Linkage groups and the physical positions are based on O_niloticus_UMD1 Assembly. The maps are color-coded: red for female specific, blue for male specific and black for sex-averaged linkage maps. Inversion in maps shows that the genetic order is inverted.

**Supplementary Figure 2:**
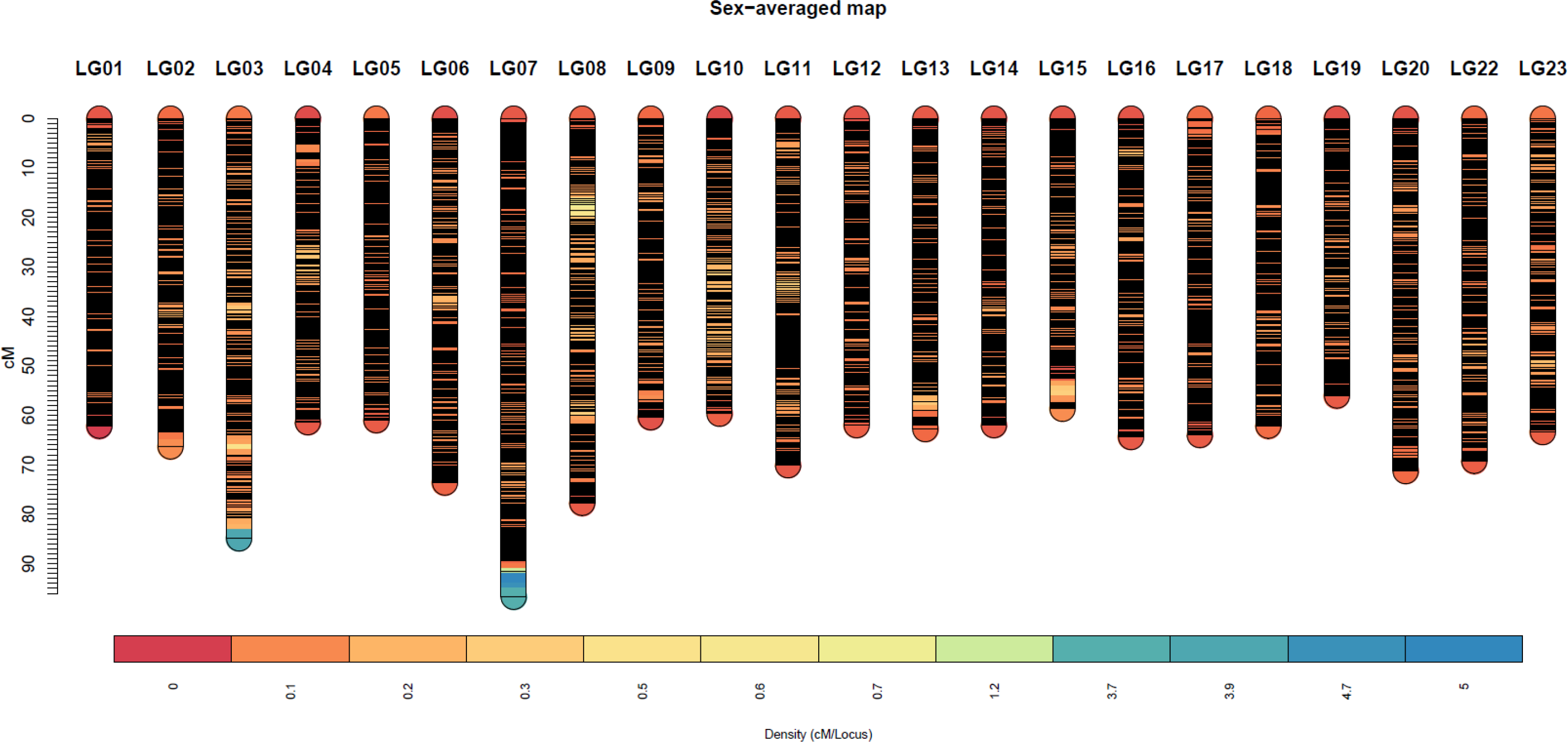
The high-density consensus (sex-averaged) Build2 linkage map for Nile tilapia. The density is measured as cM/locus (higher the value, lower the number of markers in that locus)Density (cM/Locus)

**Supplementary Figure 3:**
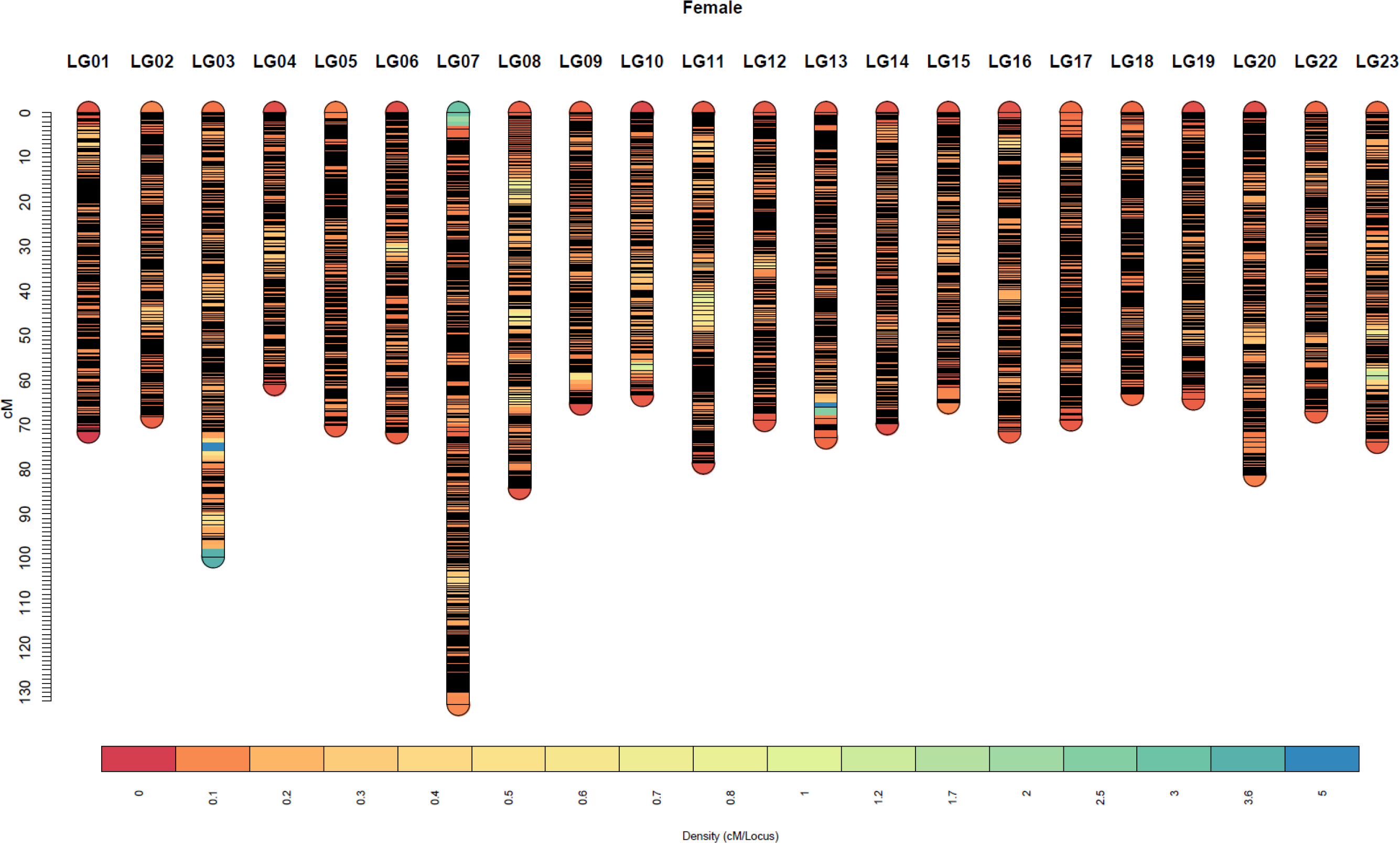
The high-density female sex specific Build2 linkage map of Nile tilapia. The density is measured as cM/locus (higher the value, lower the number of markers in that locus)

**Supplementary Figure 4:**
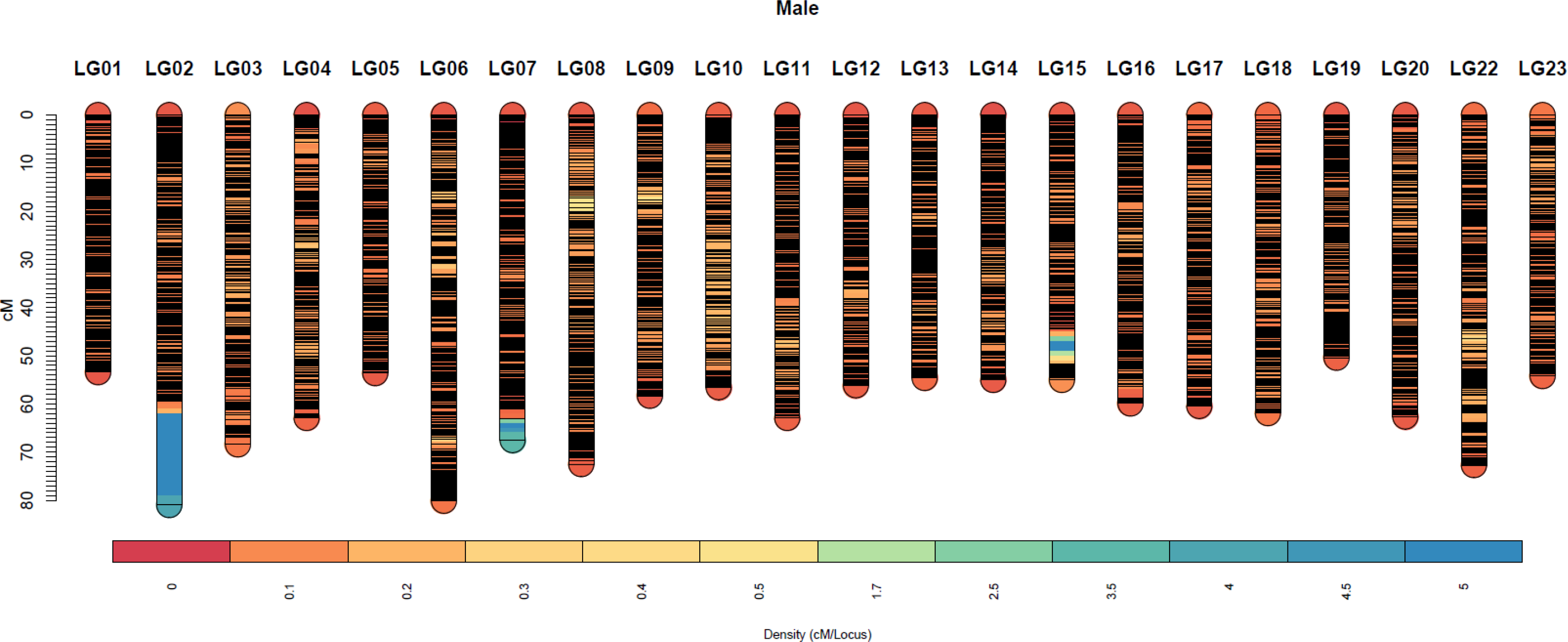
The high-density male sex specific Build2 linkage map of Nile tilapia. The density is measured as cM/locus (higher the value, lower the number of markers in that locus

